# The role of incoherent feedforward circuits in regulating precision of event timing

**DOI:** 10.1101/2020.05.17.100420

**Authors:** Supravat Dey, Sherin Kannoly, Pavol Bokes, John J Dennehy, Abhyudai Singh

## Abstract

Triggering of cellular events often relies on the level of a key gene product crossing a critical threshold. Achieving precision in event timing in spite of noisy gene expression facilitates high-fidelity functioning of diverse processes from biomolecular clocks, apoptosis and cellular differentiation. Here we investigate the role of an incoherent feedforward circuit in regulating the time taken by a bacterial virus (bacteriophage lambda) to lyse an infected *Escherichia coli* cell. Lysis timing is the result of expression and accumulation of a single lambda protein (holin) in the *E. coli* cell membrane up to a critical threshold level, which triggers the formation of membrane lesions. This easily visualized process provides a simple model system for characterizing event-timing stochasticity in single cells. Intriguingly, lambda’s lytic pathway synthesizes two functionally opposite proteins: holin and antiholin from the same mRNA in a 2:1 ratio. Antiholin sequesters holin and inhibits the formation of lethal membrane lesions, thus creating an incoherent feedforward circuit. We develop and analyze a stochastic model for this feedforward circuit that considers correlated bursty expression of holin/antiholin, and their concentrations are diluted from cellular growth. Interestingly, our analysis shows the noise in timing is minimized when both proteins are expressed at an optimal ratio, hence revealing an important regulatory role for antiholin. These results are in agreement with single cell data, where removal of antiholin results in enhanced stochasticity in lysis timing.

## 1 Introduction

Stochastic expression of gene products is an unavoidable aspect of life at the single-cell level and critically impacts functioning of cellular processes [1–14]. While the origins of stochastic expression have been extensively studied across organisms, how noisy expression of key regulatory proteins impacts timing of intracellular events is not well understood [15]. Moreover, characterization of control strategies that buffer stochasticity in event timing is critically needed to understand reliable functioning of diverse cellular processes that rely on precise temporal triggering of events.

Our prior work has used the highly malleable bacteriophage lambda as a model system for studying event timing in individual cells [16–18]. Here, an easily observable event (cell lysis) is the result of expression and accumulation of a single protein (holin) in the *E. coli* inner cell membrane up to a threshold level [19–21]. Upon reaching the critical threshold, holin nucleates to form holes in the membrane, and subsequently the cell ruptures (lyses) and phages are released into the surrounding medium. Preliminary data reveals precision in timing: lysis occurs on average at 65 min with a coefficient of variation of less than 5% [17, 18]. This precision in timing is consistent with the existence of an optimal time to lyse the infected cell [22, 23], and bacteriophage lambda may use several regulatory mechanisms to buffer random fluctuations around this optima. Intriguingly, lambda’s lytic pathway synthesizes two functionally opposite proteins: holin and antiholin from the same mRNA in a 2:1 ratio [24–26]. Antiholin sequesters holin, and prevents holin from participating in hole formation creating an incoherent feedforward circuit (Fig. 1) A key focus of this work is to characterize the role of this feedforward circuit in regulating precision in event timing.

**Fig. 1.**
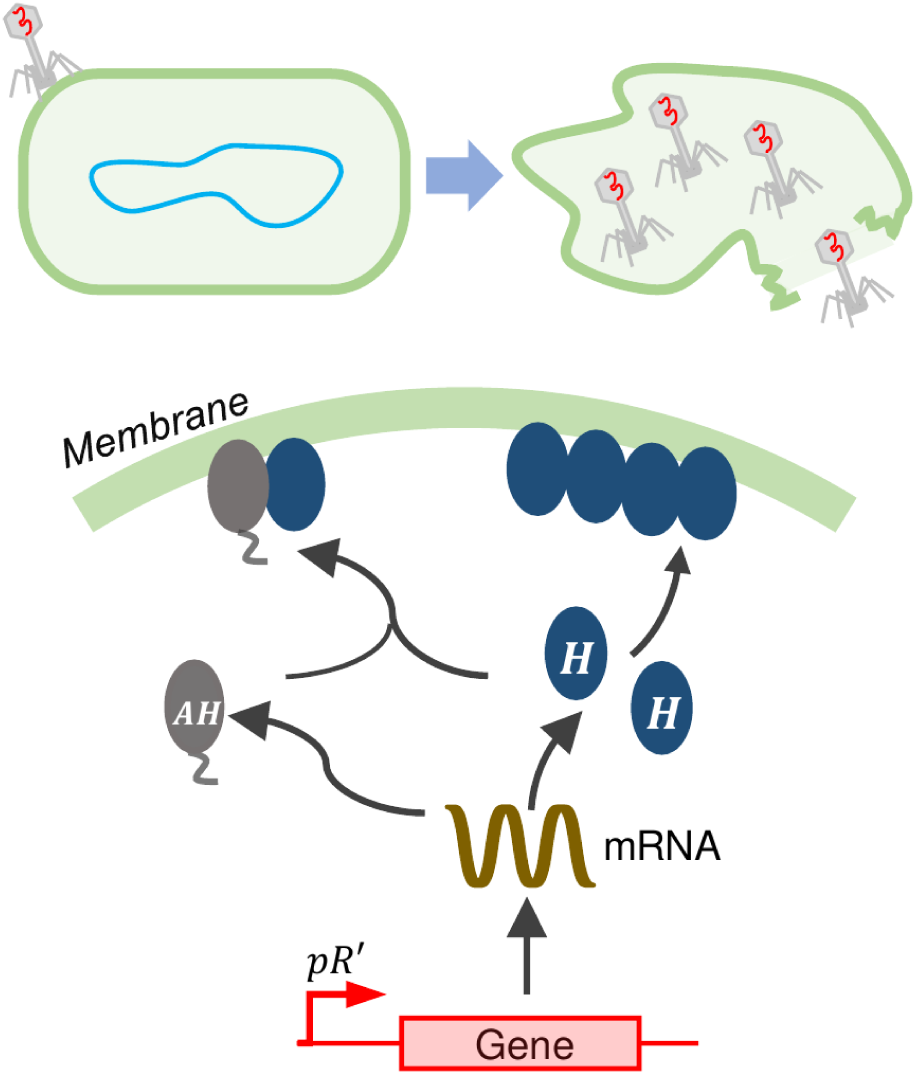
Feedforward control of phage lysis timing. *Top*: Schematic of an *E. coli* cell infected by bacteriophage lambda, and its subsequent lysis to release viral progeny. *Bottom*: After infection, the viral protein holin (H) is expressed and accumulates in the cell membrane. When holin reaches a critical threshold concentration, it forms holes and lysis ensues. The virus also encodes another protein, antiholin (AH; translated from the same mRNA transcript using a dual start site) that binds to holin and prevents it from forming holes creating an incoherent feedforward circuit.

We mechanistically model the holin-antiholin circuit using a hybrid system, where expression of both proteins occurs in stochastic bursts. The bursts arrive as per a Poisson process and result in random jumps in the protein concentrations. The binding/unbinding of antiholin to holin, and dilution of concentrations from cellular growth are modeled deterministically using mass-action kinetics. In essence, the expression of holin and antiholin in random bursts is assumed to be the predominant source of stochasticity in the feedforward circuit. As done in several recent works [27–34], we capture noise in event timing using the first-passage time framework, where lysis timing is the first time the free (unbound to antiholin) holin concentration crosses a critical threshold starting from zero initial conditions. Our analysis develops novel approximate formulas of both the mean and noise in lysis timing, and systematic analysis of these formulas elucidates the important noise-buffering role of antiholin.

### Symbols and notation

The concentrations of free holin, free antiholin and the holin-antiholin complex at time *t* inside the cell is denoted by *h*_*f*_ (*t*), *a*_*f*_ (*t*) and *c*(*t*), respectively. The total holin and antiholin concentrations are represented by *h*_*t*_(*t*) and *a*_*t*_(*t*), respectively. We use angular brackets ⟨·⟩ to represent expected values of random variables and stochastic processes, while 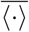 denotes the steady-state expected value.

## 2 Model Formulation

To mechanistically capture stochastic effects in the feedforward circuit, we borrow the modeling framework of bursty gene expression [35–40]. Here bursts arrive as per a Poisson process, and each burst results in the protein concentration increasing as a per burst size. In between two successive bursts, the concentrations exponentially decay due to dilution from cellular growth. Biologically, the burst arrival rate corresponds to the rate at which mRNAs are transcribed. In contrast, the burst size is the number of new proteins synthesized in a single mRNA lifespan and is determined by the mRNA translation rate. Let *h*_*f*_ (*t*) and *a*_*f*_ (*t*) denote the concentration of free (unbound) holin, and free antiholin in an individual cell at time *t*. Burst event occurs with rate *k*_*m*_, and whenever a burst occurs the concentrations jump as per the following reset map

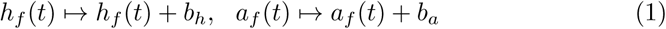

where *b*_*h*_ and *b*_*a*_ are independent and identically distributed random variables representing the burst sizes of holin and antiholin, respectively. Consistent with experimental measurements of burst sizes for *E. coli* genes [41–43], we assume *b*_*h*_ and *b*_*a*_ to follow an exponential distribution with means ⟨*b*_*h*_⟩ and ⟨*b*_*a*_⟩, respectively. Recall that for exponentially-distributed bursts, the second-order moments are given by

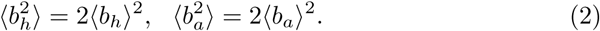

Translational of both proteins from the same mRNA transcript results in *b*_*h*_ and *b*_*a*_ being correlated. For example, a mRNA that takes a longer time to decay will translate a higher numbers of both proteins. This interdependence is characterized by the correlation coefficient *β*

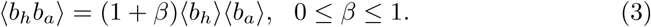

Another critical parameter of interest is

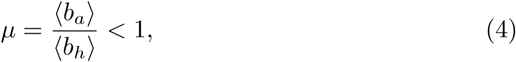

that represents the ratio of the average burst sizes as determined by the relative translation rates of these proteins from the same mRNA. Here *µ* = 0 corresponds to holin expression with no antiholin-mediated feedforward control. In the wildtype bacteriophage *λ* this ratio is reported to be *µ* = 0.5, i.e., on average one antiholin molecule is synthesized for two holin molecules [24]. We will examine in detail how stochasticity in the feedforward circuit is modulated as both *µ* and *β* are varied between 0 and 1.

To implement the feedforward circuit, free antiholin molecules bind to free holin molecules with rate *k*_*b*_ to form an inactive complex, and the complex dissociates with rate *k*_*u*_. The concentrations of holin, antiholin, and the complex (represented by *h*_*f*_, *a*_*f*_, and *c*) evolve as per the following *nonlinear* ordinary differential equations obtained using mass-action kinetics

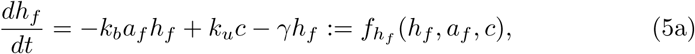

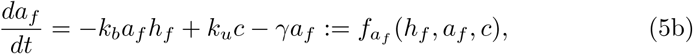

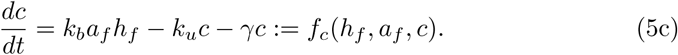

Assuming holin and antiholin are stable proteins with long half-lives, their turnover is primarily governed by dilution from cellular growth. This exponential decay in concentrations is represented by the last term in the above differential equations with *γ* being the dilution rate. In the subsequent analysis below, we will further assume that the binding/unbinding occurs sufficiently fast compared to the timescale of protein turnover, i.e., *k*_*b*_, *k*_*u*_ ≫ *γ*. In summary, we have developed a hybrid model for the holin-antiholin feedforward circuit that couples randomly occurring burst events with continuous time evolution of concentrations as per (5). Note that this model falls within the class of piecewise-deterministic Markov processes.

To characterize the stochastic dynamics of this hybrid system, we focus on the time evolution of the first- and second-order moments of *h*_*f*_ (*t*), *a*_*f*_ (*t*) and *c*(*t*). To obtain the moment dynamics we use the fact that for any arbitrary differential function 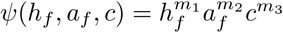 where *m*_1_, *m*_2_, *m*_3_ ∈ {0, 1, 2, …}, the time evolution of the expected value ⟨*Ψ*(*h*_*f*_, *a*_*f*_, *c*)⟩ is given by

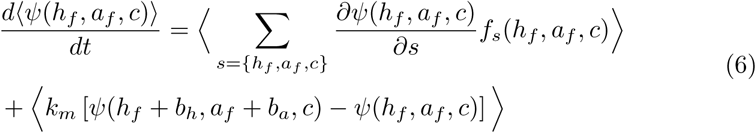

[44–46]. Thus, by appropriately choosing the positive integers *m*_1_, *m*_2_, *m*_3_ we can write the time evolution all the first- and second-order moments of *h*_*f*_ (*t*), *a*_*f*_ (*t*) and *c*(*t*). Differential equations describing the time evolution of these moments are shown in the Appendix. However, these equations cannot be solved as non-linearities in (5) result in the well-known problem of unclosed moment dynamics - the time evolution of a lower-order moment depends on higher-order moments. Generally closure schemes are employed in such cases to obtain a closed system of approximated moment dynamics [47–57]. Here we take an alternative approach based on the Linear Noise Approximation [58–61]. More specifically, assuming small fluctuations in protein levels around their respective means, the nonlinear binding term in (5) is linearized as

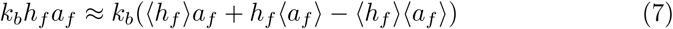

[62, 63], where ⟨*h*_*f*_⟩ and ⟨*a*_*f*_⟩ are the mean levels of the free holin, and the free antiholin, respectively. Linearizing the binding term results in closed a system of moment dynamics that can be solved to obtain both the transient and steady-state statistical moments (see Appendix). We use this approach to derive approximate formulas for the holin mean and noise levels.

## 3 Results

Having formulated a stochastic model for the feedforward circuit we characterize the noise-buffering role of antiholin in two complimentary ways:

- The effect of antiholin on steady-state fluctuations in the free holin concentration.
- The effect of antiholin on fluctuation in lysis timing, i.e., the time taken for the free holin concentration to hit a critical threshold for the first time.

We start by first investigating the extent of fluctuations in the free holin concentration.

### 3.1 Noise in the free holin level at steady-state

Let the total (free plus complex bound) holin (antiholin) concentration be denoted by *h*_*t*_ = *h*_*f*_ + *c* (*a*_*t*_ = *a*_*f*_ + *c*). Then, the steady-state average levels of total concentrations

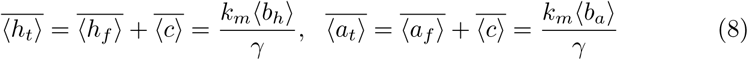

are independent of binding/unbinding rates. In the limit of fast binding/unbinding, i.e., *k*_*b*_ → ∞ and *k*_*u*_ → ∞ for a fixed dissociation constant *k*_*d*_ = *k*_*u*_*/k*_*b*_, the mean steady-state level of the free holin is given by

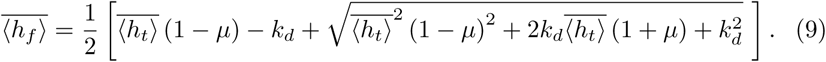

Not surprisingly, 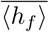 monotonically decreases with increasing amount of antiholin that is captured here by increasing the ratio *µ* of antiholin/holin burst sizes. We quantify the steady-state noise in *h*_*f*_ (*t*) using the square of the coefficient of variation (variance divided by mean squared). Our analysis yields the following formula for the noise level (see details in the Appendix)

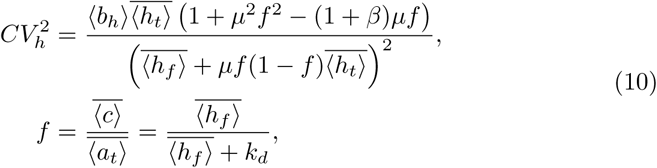

where *f* represents the fraction of total antiholin that is bound to holin. In Fig. 2(A) we plot the noise level 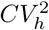 as function of *µ* for different *β* values. These results show that for a given *β*, the noise increases with increasing *µ*. However, in the limit *β* → 1 (completely correlated holin and antiholin bursts), noise decreases with increasing *µ*. It is important to point out that in this plot the mean level 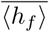 decreases with increasing *µ*.

**Fig. 2.**
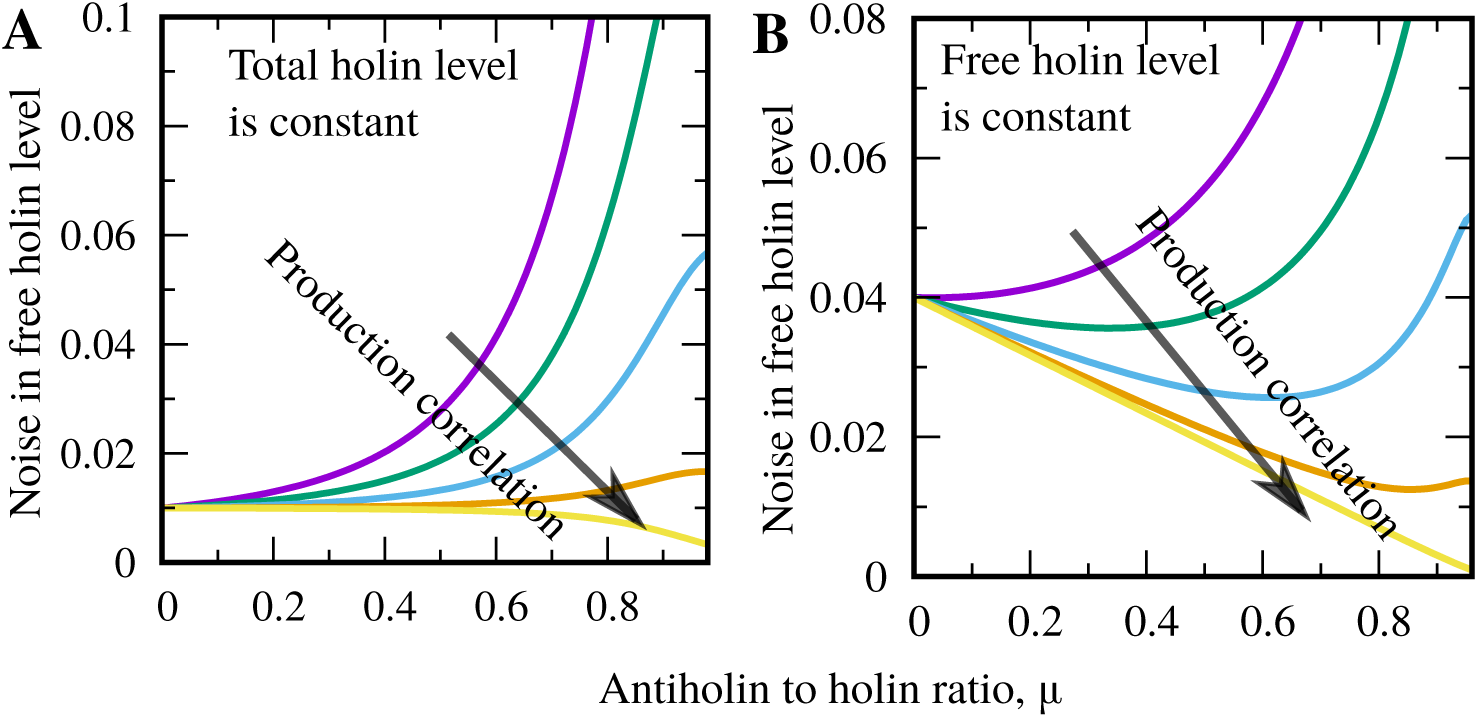
Correlated production of antiholin and holin minimizes noise in the free holin level at steady-state. (A) The noise in the free holin level decreases with increasing correlated production of holin and antiholin. The noise level 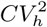 in (10) increases with *µ* for a given *β*, except when *β* → 1 where the noise decreases with *µ*. (B) The noise is plotted keeping the mean free holin level fixed 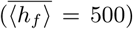 by simultaneously changing burst arrival rate *km*. In this case, for a given *β* > 0, the noise shows a minima at a critical value of *µ* = *µ*\*. The value of the minima reduces with the increment of *β* and *µ*\* → 1 as *β* → 1. Parameter used: ⟨*b*_*h*_⟩ = 10, *k*_*d*_ = 10, *k*_*m*_ = 100, *γ* = 1, *β* = 0,0.5 0.8, 0.95 and 1.

In Fig. 2(B) we plot the noise level 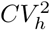 as function of *µ* by simultaneously increasing the burst arrival rate *k*_*m*_ to keep 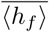 fixed. Interestingly, in this more controlled comparison, 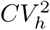 varies non-monotonically with increasing *µ* and noise is minimized at an optimal ratio *µ* = *µ*\*. To gain more analytical insight into this optimal value of *µ*, we consider the high binding affinity limit (*k*_*d*_ → 0) where *f* → 1, i.e., all the of antiholin is bound to holin. In this limit, the noise formula (10) is simplified to

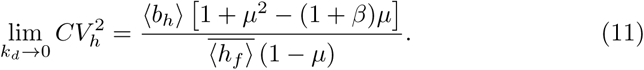

Remarkably, analysis of this simplified formula shows that holin noise level is minimized at

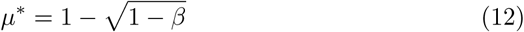

and the corresponding minimal noise is

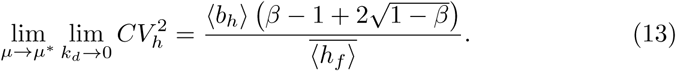

Comparing this minimal noise level with the noise level in the absence of antiholin (*µ* = 0) shows noise reduction by a factor of

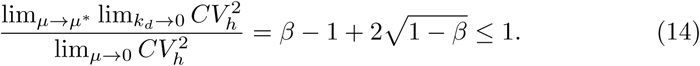

The noise attenuation only happens for *β* > 0, and becomes increasingly effective as bursts become more correlated with

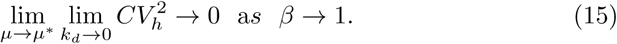

In summary, correlated expression of functionally antagonistic proteins (antiholin and holin) allows the incoherent feedforward design to effectively buffer random fluctuations in the levels of the free holin.

### 3.2 Noise in the timing lysis

We next focus on the timing of lysis that is triggered when the free holin level reaches a critical threshold for the first time. More precisely, starting from zero initial conditions, the lysis time can be mathematically formulated as the first-passage time

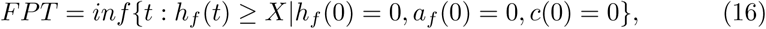

where *X* is the critical threshold level of free holin needed for lysis to occur. For the sake of simplicity, we will quantify fluctuations in *FPT* only in the limit of high binding affinity (*k*_*d*_ → 0) where all the antiholin is bound to holin, and the amount of free holin can be approximated as

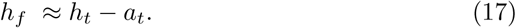

Using unclosed moment dynamics for the free holin, free antiholin, and complex concentrations (presented in the Appendix), the time evolution of the statistical moments of the total holin (*h*_*t*_ = *h*_*f*_ + *c*), and the total antiholin (*a*_*t*_ = *a*_*f*_ + *c*) concentrations are given by

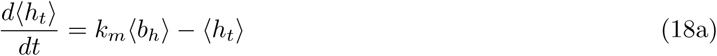

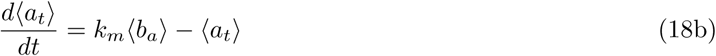

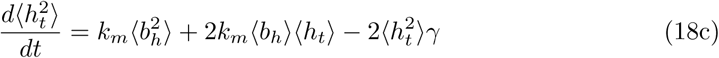

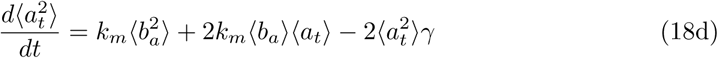

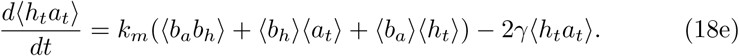

An advantage in working with the total concentrations is that their moment dynamics can be solved exactly from (18) as the nonlinear binding terms are absent here. The transient and steady-state moments of the free holin level can then be obtained using the approximation (17). From (18), the time evolution of the mean levels ⟨*h*_*t*_⟩ and ⟨*a*_*t*_⟩

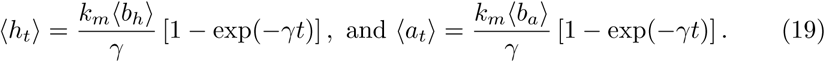

shows that the ratio of total holin to antiholin remans invariant over time, i.e. ⟨*a*_*t*_⟩*/* ⟨*h*_*t*_⟩ = ⟨*b*_*a*_⟩*/* ⟨*b*_*h*_⟩ = *µ*. From (17), the mean free holin level evolves as

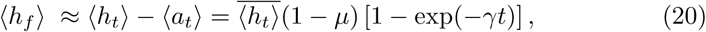

and assuming small concentration fluctuations, the average time taken by *h*_*f*_ to reach the threshold *X* will be

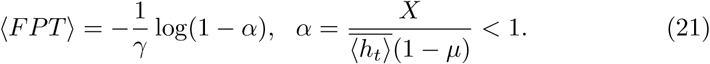

Here *α* < 1 can be interpreted as the ratio of the critical threshold needed for lysis to the steady-state free holin level. As expected, the average lysis time ⟨*FPT*⟩ increases with increasing levels of antiholin that sequesters the functionally active holin (Fig. 3(A)).

**Fig. 3.**
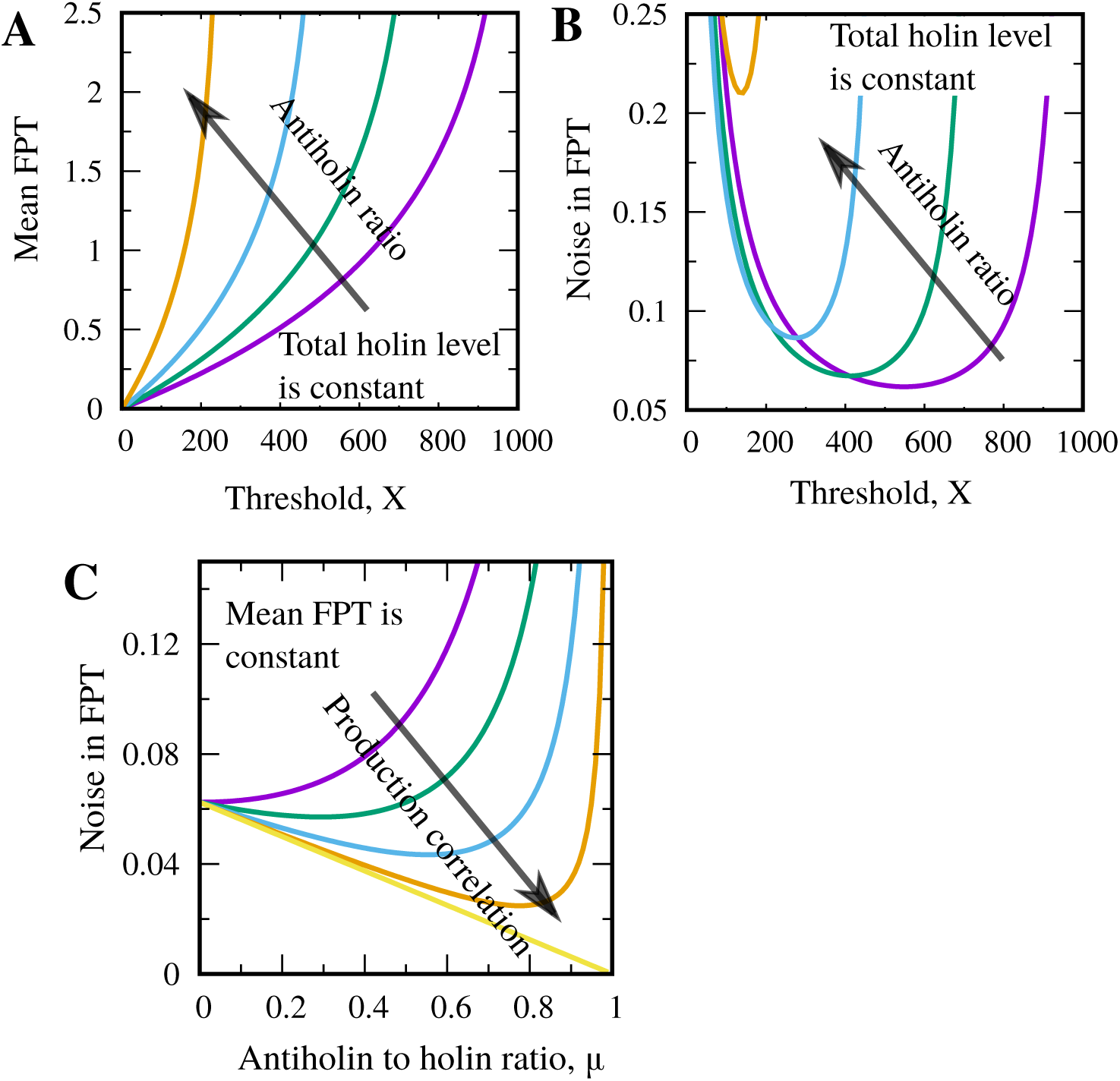
Correlated production of antiholin and holin reduces the noise in timing. (A) The mean lysis time ⟨*FPT*⟩ against threshold level *X* for different antiholin-to-holin ratios (*µ* = 0, 0.25, 0.5, and 0.75). The presence of antiholin increases ⟨*FPT*⟩ for a given *X*. (B) The noise in lysis timing 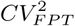 is plotted as a function of the threshold level *X* for different ratios (*µ* = 0, 0.25, 0.5, and 0.75) with *β* = 0.8. The timing noise is minimized at an optimal threshold level. (C) The noise in lysis timing 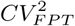 as a function of *µ* with a fixed mean ⟨*FPT*⟩ = 0.693 *γ*^*−*1^ for different values of *β* = 0, 0.5, 0.8, 0.95, and 1. The noise decreases with enhanced correlation between holin and antiholin production. We increase *km* to keep ⟨*FPT*⟩ fixed and other parameters are taken as: ⟨*b*_*h*_⟩ = 10, *k*_*m*_ = 100, and *γ* = 1.

Next we turn our focus on the noise in *FPT*, which in the limit of small fluctuations can be approximately as

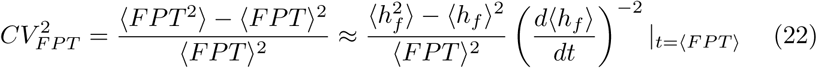

[27]. In essence, the noise in FPT (as measured by the square of the coefficient of variation) is determined by the noise in the free holin level at *t* = ⟨*FPT*⟩ and the slope (*d* ⟨*h*_*f*_⟩ */dt*) at which ⟨*h*_*f*_⟩ hits the threshold *X*, with shallower slopes amplifying the noise in *FPT* [27]. From (17) and (18), the transient variance in the level of the free holin is obtained as

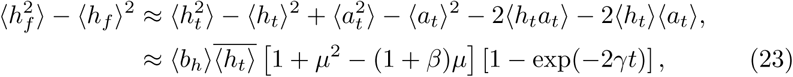

which using (20) and (22), yields the following noise in lysis timing

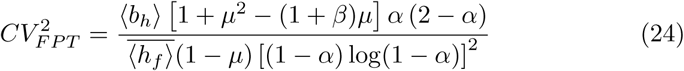

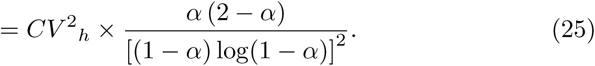

Here *CV* ^2^_*h*_ is the steady-state noise in the free holin level in the high binding affinity limit as determined in (13), and *α* is the ratio of the threshold *X* to 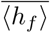.

Analysis of the noise in timing formula (25) reveals two intriguing results. Firstly, 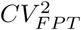 varies non-monotonically with increasing *α* (and hence, increasing threshold *X*) and is minimized at an optimal threshold (see Fig. 3(B))

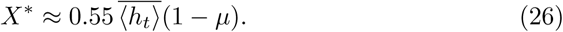

Intuitively, noise in timing is high at a low threshold due to the low number of burst events taken to cross threshold. Noise is also amplified at a high threshold as the slope (*d*⟨*h*_*f*_⟩/*dt*) becomes shallower, resulting in the highest timing precision at an intermediate threshold (26). A second insight from (25) is that for a fixed mean lysis timing, noise in timing is minimized at the optimal ratio *µ* = *µ*\* given by (13) when *β* > 0 (Fig. 3(C)), and this effect arises straightforwardly from the fact that *CV* ^2^_*h*_ is minimized at *µ* = *µ*\*.

## 4 Discussion

We have systematically investigated the role of an incoherent feedforward circuit in regulating timing of a key bacteriophage life history event - the lysis of the infected host cell to release phage progeny. After infecting an *E. coli*. cell, phage lambda makes a key developmental decision between lysis and lysogeny [64–67]. If the lysis pathway is chosen, then a suite of lysis proteins (that includes holin and antiholin) are constitutively expressed from the lambda genome. Holin and antiholin accumulate in the *E. coli* inner membrane with antiholin sequestering holin into an inactive complex [25]. Lysis occurs when the free holin concentration in the membrane crosses a critical threshold that triggers holin nucleation and hole formation [68]

To understand the noise-buffering role of antiholin, we developed and analyzed a stochastic model of the feedforward circuit. Our analysis shows that for fixed mean levels, the steady-state noise in the free holin concentration (Fig. 2), and the timing of lysis (Fig. 2), are minimized when the antiholin-to-holin production ratio is optimally set at *µ* = *µ*\* given by (13). The corresponding noise formulas reveal the limits of noise reduction reached by a feedforward circuit as compared to an open-loop circuit in the absence of antiholin. These results are consistent with other computational and experimental studies illustrating the noise-suppression ability of incoherent feedforward circuits [69–74] that have primarily focused on noise in protein levels. The novelty of this work stems from considering correlated expression and dimerization of two antagonistic proteins, and quantifying the impact of this interaction on the statistics of event timing.

Our modeling results are consistent with single-cell measurements where the wild-type bacteriophage lambda encoding both holin and antiholin has reduced stochasticity in lysis timing as compared to a mutant lambda that lacks antiholin [17,18]. Finally, results presented here provide rich predictions that can be tested with further experiments. The dual start-site mRNA transcript that synthesis holin and antiholin has a hairpin loop just upstream of the translational start site. This hairpin loop regulates ribosomal access to the start sites, and hence sets the ratio *µ* [24]. Mutations in the hairpin loop can be used to alter *µ*, providing an exciting model system to probe the role of feedforward circuits in regulating precision of event timing.

## Appendix

### Moment dynamics for the free holin, free antiholin and complex concentrations

Using (6) in the main text, we obtain moment dynamics for the first and second-order in *h*_*f*_, *a*_*f*_, and *c*:

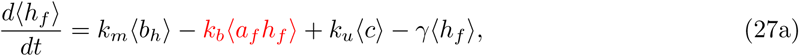

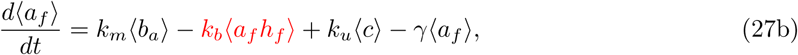

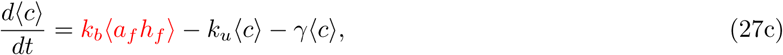

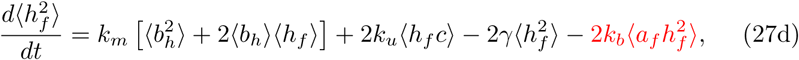

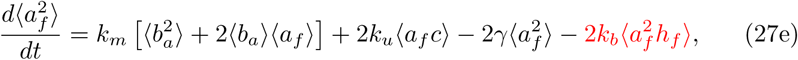

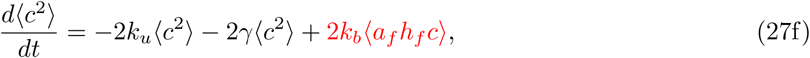

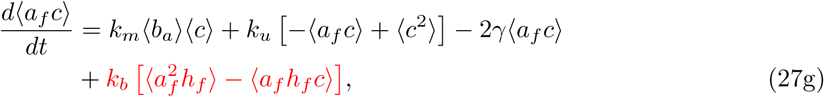

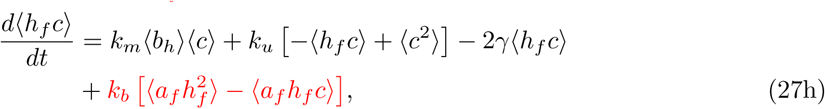

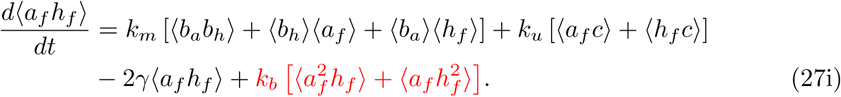

The above dynamical equations for moments are unclosed due to the presence of higher-order moments (marked in red). We assume small fluctuations around the mean to linearize the binding term as: *k*_*b*_*h*_*f*_ *a*_*f*_ = *k*_*b*_(⟨*h*_*f*_⟩ *a*_*f*_ + *h*_*f*_ ⟨*a*_*f*_⟩ *−* ⟨*h*_*f*_⟩ ⟨*a*_*f*_⟩), and derive a closed system of moment dynamics given by

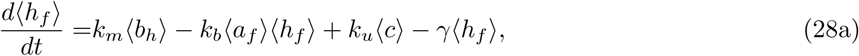

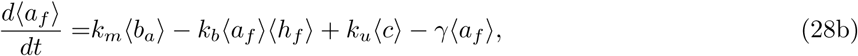

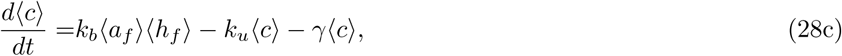

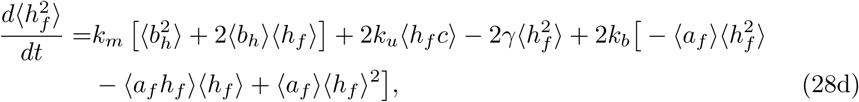

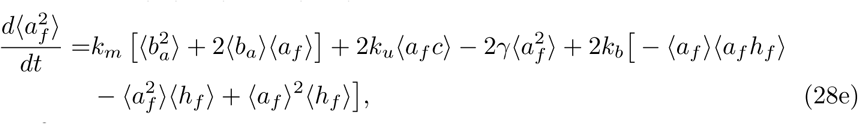

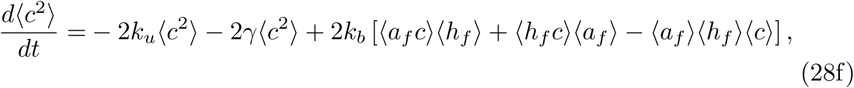

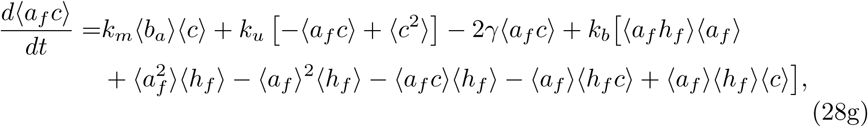

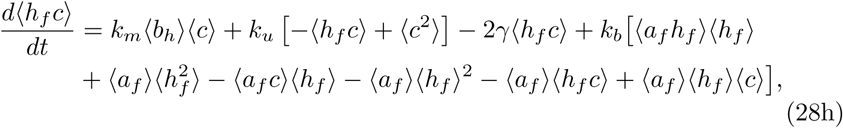

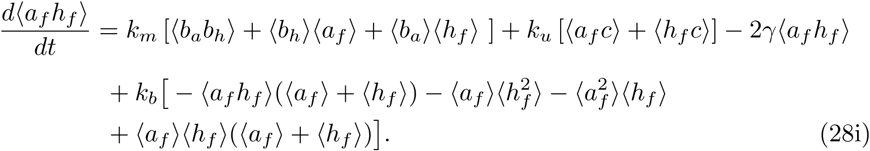

We use the above closed moment equations to calculate the formulas for the mean and noise in the free holin level at steady state. The analytical expression presented in the main text are obtained in the fast binding/unbinding limit i.e., *k*_*b*_ → ∞ and *k*_*u*_ → ∞ for fixed dissociation constant *k*_*d*_ = *k*_*u*_*/k*_*b*_.

## Acknowledgement

AS was supported by NIH grants 5R01GM124446 and 5R01GM126557. PB was supported by the Slovak Research and Development Agency under the contract No. APVV-18-0308, by the VEGA grant 1/0347/18, and the EraCoSysMed project 4D-Healing. SD would like to thank Khem Raj Ghusinga for insightful discussions.

